# BMSC-Derived Exosomal MiRNAs Can Modulate Bone Restoration in Diabetic Rats with Femoral Defects

**DOI:** 10.1101/2021.09.07.459238

**Authors:** Ning Wang, Xuanchen Liu, Zhen Tang, Xinghui Wei, Hui Dong, Yichao Liu, Hao Wu, Zhigang Wu, Xiaokang Li, Xue Ma, Zheng Guo

## Abstract

The exosomal miRNAs of BMSCs participate in hyperglycemia induced poor healing of bone defects. Here, we demonstrate that exosomes derived from BMSCs harvested from diabetes mellitus(DM) rats suppressed bone formation when administered to normal rats with bone defects. Using high-throughput sequencing analysis of microRNA molecules, high miR-140-3p levels were expressed in exosomes released by N-BMSCs. Using TargetScan software and luciferase activity assays, *plxnb1* was identified as the downstream molecular target of exosomal miR-140-3p that regulated osteogenesis. Transplantation of exosomes that overexpressed miR-140-3p into DM rats promoted the restoration of bone defects. Furthermore, miR-140-3p significantly promoted the differentiation of DM-BMSCs into osteoblasts and inhibited the expression p-RohA and p-ROCK in the plexin B1 signaling pathway. Taken together, these observations suggest that DM decreases the levels of exosomal miR-140-3p, which impedes bone formation and the differentiation of BMSCs. MiR-140-3p may represent a potential therapeutic target for DM related to abnormal bone metabolism.

## Introduction

Type 2 diabetes mellitus (T2DM) is a common metabolic disorder characterized by chronic hyperglycemia. According to the World Health Organization (WHO), more than 463 million people around the world are currently suffering from diabetes mellitus (DM), of which T2DM accounts for 90% (Rauch *et al*, 2019). There is accumulating evidence of a strong interaction between glucose levels and changes in bone metabolism. Impaired bone healing is considered an important complication associated with DM. T2DM also results in changes in bone metabolism detrimental to bone quality and a cause of reduced bone strength, increased fracture risk, and impaired bone healing (Compston, 2018; Jiao *et al*, 2015; Paschou *et al*, 2017). It has been reported that the duration of fracture healing in DM patients is prolonged by 87% (Hu *et al*, 2019). Bone regeneration is often delayed in DM patients, most likely due to impaired osteogenic differentiation (Gopalakrishnan *et al*, 2006; Hamann *et al*, 2011). Furthermore, the inhibition of transcription factor expression, which is crucial for the development of an osteoblastic phenotype such as runt-related transcription factor 2 (Runx-2), and decreased cell proliferation and growth factor expression contribute to impaired bone healing (Marin *et al*, 2018). Despite such evidence, the precise mechanisms causing bone impairment in T2DM are not yet fully understood. Further investigation is required to elucidate the pathophysiology of diabetic bone and develop successful strategies to treat this growing medical complication that has led to considerable socioeconomic burden.

Bone mesenchymal stem cells (BMSCs), also known as bone marrow-derived mesenchymal stem cells (MSCs), are considered a promising source of cells for tissue engineering because they can stimulate osteogenesis required for bone regeneration (Garcia-Garcia *et al*, 2015; Gomez-Barrena *et al*, 2019). Recent evidence indicates that the osteogenic capability of BMSCs is reduced in DM patients in comparison with that in normal populations (Guo *et al*, 2017). The ability of BMSCs to self-renew and their therapeutic potential play important roles in regulating the healing of bone defects (Gomez-Barrena *et al*., 2019). Recently, multiple studies have revealed that the efficacy of MSC-based therapies for bone defect regeneration is attributable to their exosome-mediated autocrine and paracrine actions (Bjorge *et al*, 2017; Kourembanas, 2015; Toh *et al*, 2017; Zhu *et al*, 2019). MSC-derived exosomes are more stable for therapeutic interventions in particular physiological circumstances due to the additional immune privilege compared with stem cells (Kishore & Khan, 2017; Liu *et al*, 2017b). Extensive studies have demonstrated that BMSC-derived exosomes can stimulate the proliferation and osteogenic differentiation of BMSCs, thereby promoting osteogenesis, angiogenesis, and bone mineralization within bone defects (Qin *et al*, 2016; Sahoo *et al*, 2011), suggesting that they could have potential as a cell-free therapy for bone defect healing.

Exosomes are extracellular vesicles, ranging in diameter from 30 to 150 nm and thought to be the primary mediators of intercellular communication, by means of small RNA molecules, especially microRNAs (miRNAs) (Ameres *et al*, 2007; Bartel, 2004; Brennecke *et al*, 2005; Jeppesen *et al*, 2019). A number of bone-derived exosomal miRNAs that regulate bone remodeling have been characterized. Exosomes are found in the circulation and contain numerous miRNA molecules which perform important regulatory roles at both local and distal sites. It has been revealed that BMSC-derived exosomes promote osteoblastogenesis via miR-196a *in vitro* and have been shown to increase bone regeneration in calvarial defects in a Sprague-Dawley rat model (Qin *et al*., 2016). A previous study found that osteoclast-derived exosomal miR-214-3p can be transferred to osteoblasts, inhibiting osteogenic activity and bone formation. However, inhibition of miR-214-3p expression in osteoclasts promoted osteogenesis and bone regeneration (Li *et al*, 2016). Moreover, miR-31a-5p expression increased significantly in BMSCs and BMSCs-derived exosomes from elderly rats. However, BMSCs in which miR-31a-5p was overexpressed displayed increased adipogenesis and decreased osteogenesis. Inhibition of miR-31a-5p prevented substantial bone loss and decreased osteoclastic activity in elderly rats (Xu *et al*, 2018). In addition, miR-100-5p derived from MSCs harvested from the infrapatellar fat pad (IPFP) was found to reduce autophagy in chondrocytes to a significant extent by inhibition of the mTOR autophagy pathway, protecting OA mice and normalizing cartilage homeostasis (Wu *et al*, 2019).

The results described above suggest that DM may inhibit BMSC differentiation by modification of miRNA expression, which modulates osteogenesis. In the present study, we found that exosomes secreted by BMSCs derived from type 2 diabetic rats (DM-Exos) delayed the healing of bone defects in normal rats while those from normal rats (N-Exos) promoted bone formation in diabetic rats. In addition, DM-Exos reduced levels of mineral deposits while N-Exos increased mineralization by BMSCs *in vitro*. The expression of miR-140-3p in exosomes derived from type 2 diabetic rat BMSCs was lower than that in normal rat BMSCs when analyzed by high-throughput sequencing. Furthermore, plxnb1 was found using TargetScan software to be the downstream target of exosomal miR-140-3p. Notably, miR-140-3p overexpression or inhibition modulated plxnb1 expression and significantly enhanced BMSC differentiation or impeded BMSC osteoblastogenesis. Thus, a mixture of matrigel and exosomes overexpressing miR-140-3p significantly enhanced bone formation compared with N-Exos and DM-Exos treatment groups. Taken together, the present study revealed a role for BMSC-derived exosomal miR140-3p in the regulation of DM-associated impaired bone healing, suggesting that plexin B1 may be a promising target for miR-140-3p.

## Results

### BMSCs secrete exosomal miRNAs

Firstly, BMSC-derived exosomes were isolated, their contents identified, and their morphology assessed by the expression of specific markers and by TEM, respectively. The exosomes were isolated from BMSCs (CD29^+^Sca-1^+^) from the bone marrow cells of 3-month-old normal rats (N-rats) and diabetic rats (DM-rats). The two groups of BMSCs were then cultured in exosome-free medium for 72 h, from which conditioned medium was collected, then dead cells and debris were removed. Ultracentrifugation was used to isolate the BMSC-derived extracellular particles. The morphology of the isolated extracellular particles was observed by TEM, revealing that they were round, lipid-bilayered vesicles (Fig 1A), confirming that they were exosomes. A nanoparticle tracking system was used to measure the concentration and size distribution of the purified particles. There were 5.4×10^6^ with a mean diameter of 133 nm (Fig 1B). The expression of the exosome-specific protein markers HSP70, TSG101, CD63, and CD9 identified by Western blot analysis also demonstrated that the extracellular particles isolated from N-rats and DM-rats were exosomes (Fig 1C). In addition, there were no differences between N-rats and DM-rats in the expression of the exosome-associated protein markers. The results above suggest that the BMSC-derived extracellular particles were exosomes.

**Fig 1.**
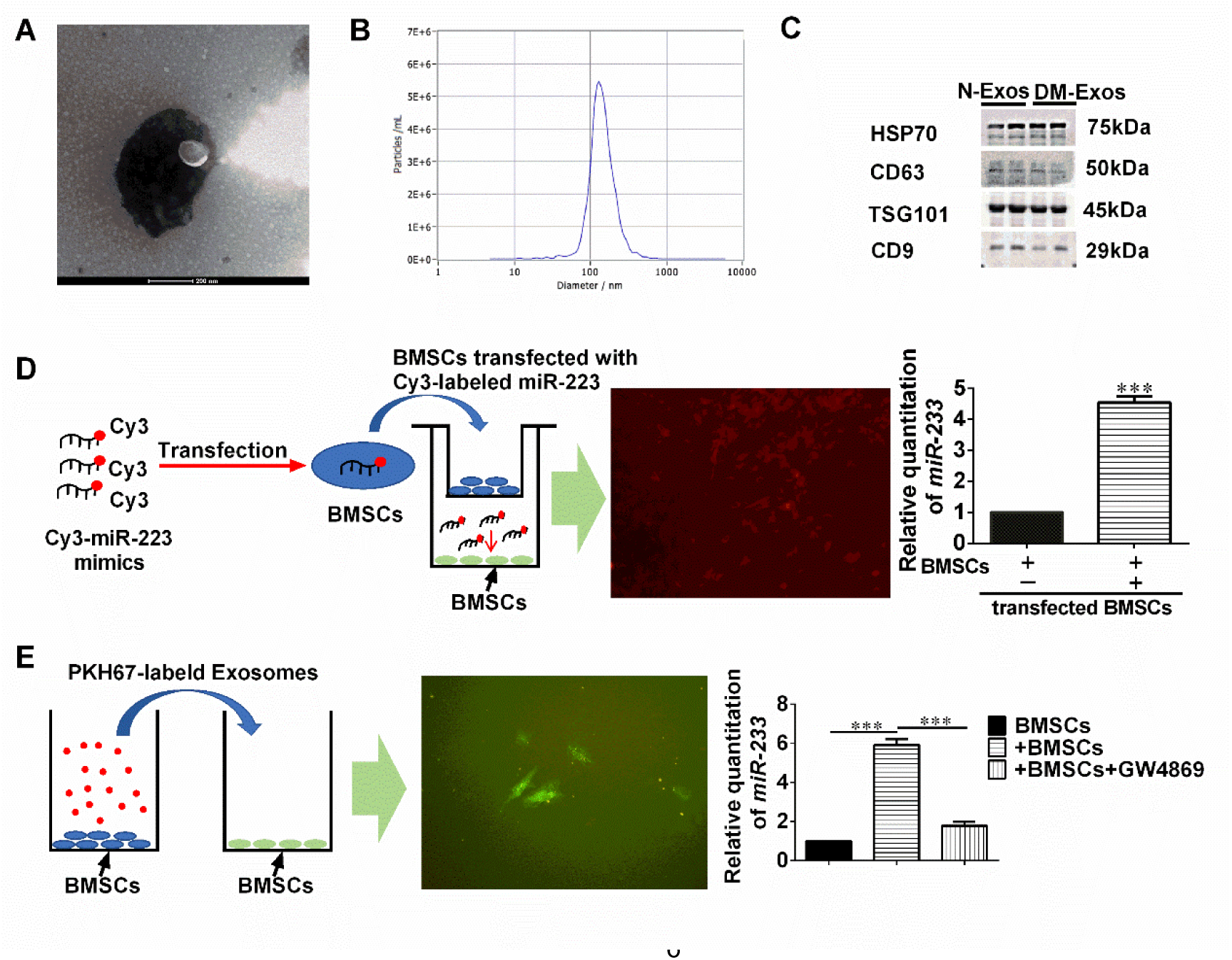
**BMSCs Secrete Exosomal miRNAs** (A) Electron microscopic analysis of Exos secreted by BMSCs. Scale bar: 200 nm. (B) Particle size distribution of vesicles secreted from BMSCs, as measured by NanoSight analysis. (C) The concentrations of the exosome (Exo)-specific marker TSG101 and extracellular vesicle-related protein markers HSP70, CD63, and CD9 were measured by Western blot analysis. The blots are representative of 3 replicates of independent experiments, each with 2 samples. (D) BMSCs transfected with a Cy3-labeled miR-223 mimic were co-cultured with BMSCs in a transwell (membrane pore size: 0.4 μm) plate. (E) Exos from BMSCs were labeled with PKH26 and then added to BMSC cultures. The results demonstrate the effects of the EV secretion inhibitor GW4689 (10 mM) on exosome-dependent miRNA delivery from BMSCs into recipient BMSCs. Data represent means ± SD. ***p < 0.001.

Whether BMSCs could secrete extracellular miRNAs able to be phagocytized by target cells was subsequently determined. MiR-223 is specific for myeloid cells and so was used as a marker. BMSCs were transfected with fluorescent Cy3-labeled miR-223 mimics then co-cultured with untransfected BMSCs seeded in the upper and lower wells of a transwell plate, respectively, for 12 h (Fig 1D). Red fluorescent Cy3 dye in the untransfected BMSCs demonstrated that the Cy3-miR-223 mimic was transferred from the BMSCs in the upper well to those in the lower well (Fig 1D). The quantities of miR-223 in the BMSCs in the lower well increased by almost 5-fold following co-culture. In addition, BMSCs were also cultured with Cy3 alone (without miR-223), and little red fluorescence was observed in these cells (Cy3-BMSCs), while no fluorescence was observed in BMSCs seeded in the lower well when co-cultured with the Cy3-BMSCs (Fig EV1). Taken together, the results indicate that BMSCs can secrete extracellular miRNAs that can be absorbed by other cells such as BMSCs.

Finally, whether BMSC-derived exosomes could be absorbed by other BMSCs was determined. BMSCs-derived exosomes were labeled with the fluorescent dye PKH67 and then added to a fresh culture of BMSCs. After 12 h, the BMSCs fluoresced green, indicating that the exosomes were absorbed by the BMSCs (Fig 1E). In addition, a transwell experiment was also performed to verify whether BMSCs can secret miRNA-containing exosomes to other cells. BMSCs seeded in the lower well expressed an approximately 6-fold increase in miR-223 when co-cultured with BMSCs seeded in the upper well. However, the extracellular vesicle secretion inhibitor GW4869 reversed the increase in miR-223 expression in BMSCs seeded in the lower well when co-cultured with BMSCs seeded in the upper well (Fig 1E). All the experiments demonstrated that BMSCs secrete miRNAs containing exosomes that could be delivered to target cells.

### Differential effects of exosomes derived from DM and normal rats on the restoration of bone defects

BMSCs play a vital role in bone remodeling. There is considerable evidence indicating that the ability of BMSCs to differentiate into osteoblasts declines in patients with diabetes compared with those without. Therefore, the ability of exosomes secreted by DM-BMSCs and N-BMSCs to modulate bone formation *in vivo* was tested.

Both normal and diabetic rats with bone defects received N-Exos(240 μg) or DM-Exos (240 μg) derived from normal rat BMSCs or DM rat BMSCs (Fig 2A). Micro-CT was used to quantify the bone mass within the defect in each group (Fig 2B,C). The results demonstrate that although treatment with exosomes increased bone mass, N-Exos caused the formation of a greater quantity of bone than DM-Exos after 2 and 8 weeks, as shown by the difference in BV/TV values. In defects receiving DM-Exos, the BV/TV in 8 weeks was higher than that in 2 weeks. Similarly, trabecular number and trabecular thickness also increased due to the transplantation of exosomes compared with the control group, while treatment with N-Exos increased trabecular number and thickness compared with rats treated with DM-Exos. Furthermore, the trabecular number and trabecular thickness after 8 weeks were also higher than after 2 weeks in the DM-Exos group. The trabecular spacing decreased in the exosome-treated groups compared with the control group. The data above indicate that although exosomes promoted bone formation, DM-Exos impaired bone formation compared with the N-Exos treatment in both normal rats and those with diabetes mellitus.

**Fig 2.**
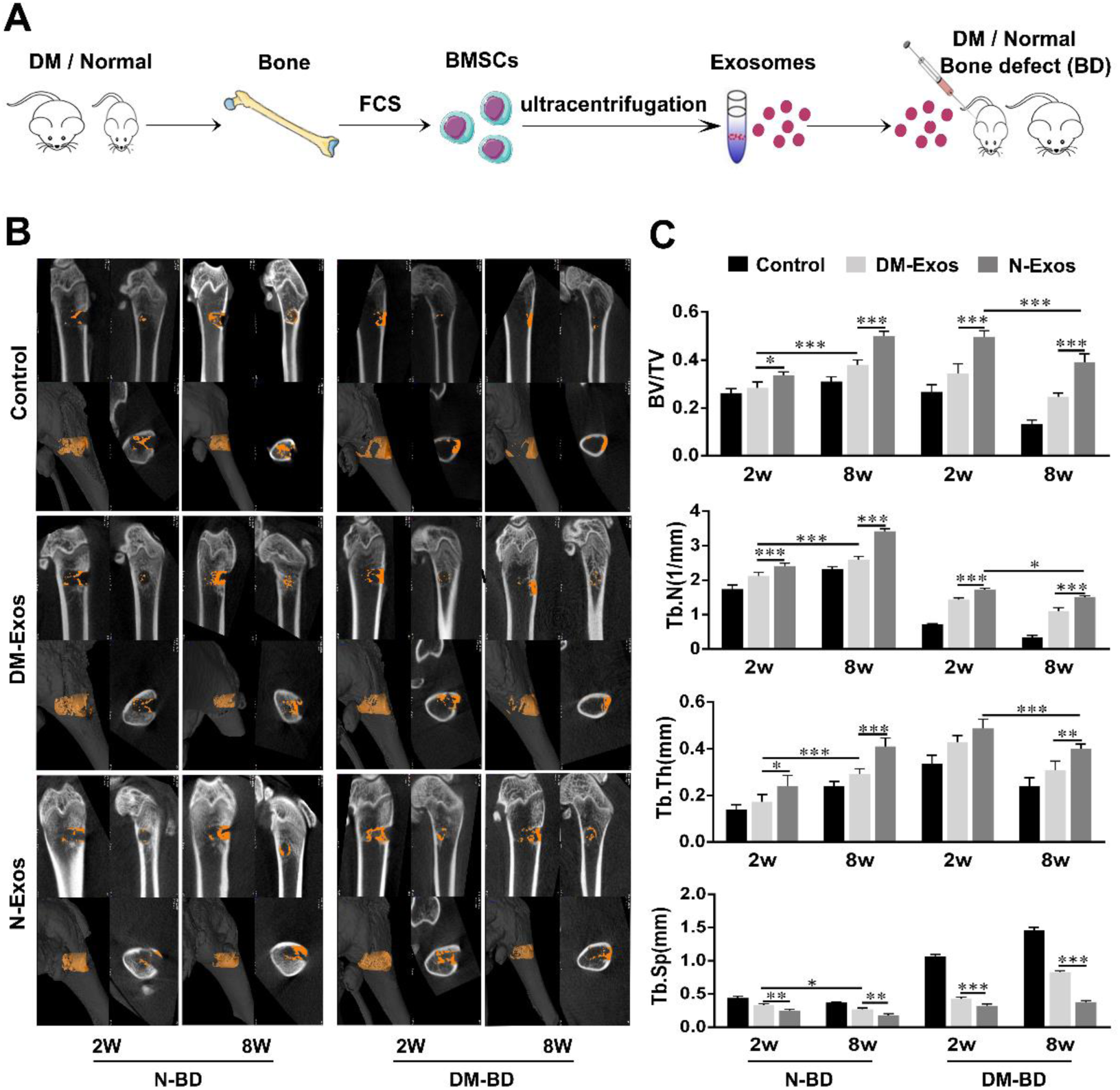
**Impaired bone formation in rat bone defect caused by DM-Exos** (A) N-Exos or DM-Exos were implanted into bone defects of rats that were normal or suffered diabetes mellitus. (B) Representative micro-CT images within a region of interest (ROI) representing bone defects 2 and 8 weeks after surgery in normal rats and those with diabetes mellitus. n=5 in each group. (C) Quantitative analysis of micro-CT imgaes of the new bone formation in bone defects 2 and 8 weeks after transplantation with N-Exos and DM-Exos. BV: bone volume; TV: tissue volume; Tb,N: trabecular number; Tb.Th: trabecular thickness; Tb.Sp: trabecular Spacing. n=5 in each group; *p < 0.05; **p < 0.01; ***p < 0.001. Data represent means ± SD.

### Effect of N-Exos and DM-Exos on bone regeneration in a rat bone defect model

Histological analysis and immunohistochemical staining were utilized to demonstrate the different effects of exosomes isolated from DM and normal rats on bone regeneration and mineralization. Collagen is a constituent of mineralized bone. In hematoxylin and eosin (HE)-stained sections (Fig 3A), collagen can be visualized as light pink tissue. In the bone defect of normal rats, little mineralized bone tissue was observed in the DM-Exos transplanted group in comparison with the N-Exos treatment group, although more than in the control group. The quantity of collagen observed in the bone defect in DM rats was consistent with that in the defects of normal rats. The N-Exos transplanted group exhibited a greater quantity of mineralized bone tissue than the DM-Exos treatment group.

**Fig 3.**
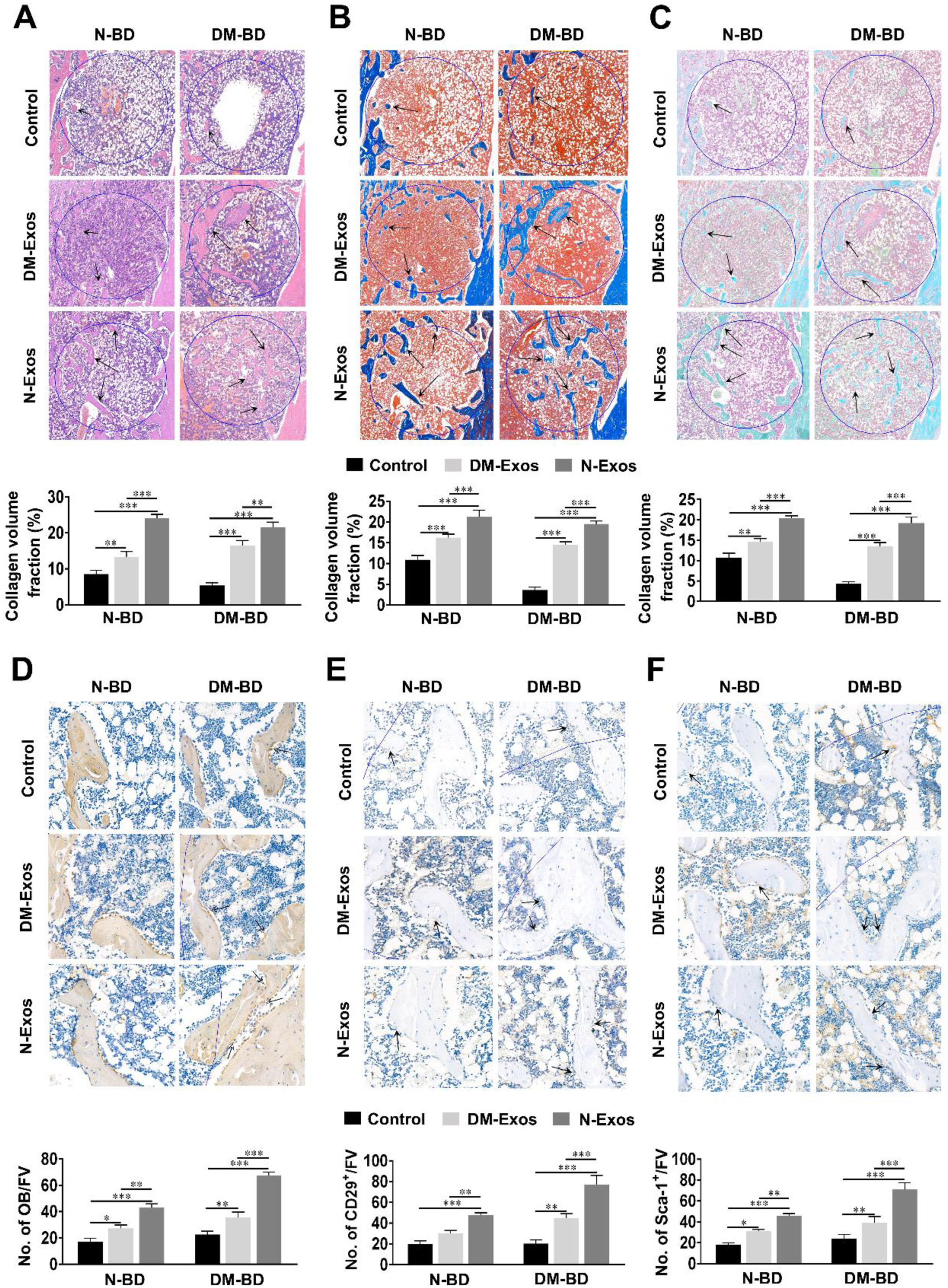
**Histological/Immunohistological analysis of bone formation after transplantation of exosomes.** (A-C) Histological analysis of decalcified sections in both normal and diabetic rats. HE/Masson’s/ Safranin O-fast green staining of new bone formation in the bone defect. The black arrow indicates new bone in light pink, blue, or green within the circle, indicating the 3mm-diameter defect. New bone formation in the bone defects was analyzed using ImageJ software. n=5 for each group. (D-F) Immunohistological analysis of the osteoblasts and BMSCs in both normal and diabetic rats. Collagen-1 and CD29/Sca-1 staining of osteoblasts and BMSCs in bone defects. Black arrows indicate osteoblasts and BMSCs on the bone surface. n=5 in each group; *p < 0.05; **p < 0.01; ***p < 0.001. Data represent means ± SD.

Collagen in bone is visualized as blue tissue in Masson’s trichrome-stained sections (Fig 3B). In normal rats with bone defects, the volume of mineralized bone tissue in the DM-Exos transplanted group was less than in the N-Exos group. The volume of mineralized bone tissue in untreated bone defect in normal rats was less than that in the exosome-treated group. Consistent with the results of bone defects in DM rats, the volume of mineralized bone tissue in the N-Exos transplanted group was greater than that observed in the DM-Exos group. In addition, the volume of mineralized bone tissue in the control group was less than in the Exos-treated groups. In Safranin O/fast-green stained sections (Fig 3C), the collagen was identified as being green. For bone defects in normal rats, the volume of mineralized bone stained green was lower in the DM-Exos group than in the N-Exos group. The quantity of mineralized bone was least in normal rats groups. Furthermore, the volume of mineralized bone in the bone defects of DM rats was lower in the DM-Exos group than in the N-Exos group. Moreover, the mineralized bone volume of the control group was the smallest of the comparative groups. Immunohistochemical staining of collagen-1 (Fig 3D) stained osteoblasts. Normal rats with bone defects had fewer osteoblasts in the control group than in either the DM-Exos or N-Exos group, while the numbers in the N-Exos group were higher than in the DM-Exos group. The numbers of osteoblasts in the bone defects of DM rats were smaller in the control group than in either the DM-Exos or N-Exos group, while there were more osteoblasts in the N-Exos group than in the DM-Exos group. BMSCs can differentiate into osteoblasts, losing the CD29/Sca-1 markers specific for BMSCs. In the bone defects of normal rats (Fig 3E,F), the number of BMSCs on the bone surface in the DM-Exos treatment group was less than in the N-Exos group, which was more than in the control group. The numbers of BMSCs on the bone surface in the N-Exos treatment group in DM rats was considerably greater than in the DM-Exos treatment group, which was more than in the control group. The data above indicate that N-Exos promoted bone formation via increasing the numbers of BMSCs and osteoblasts in DM rats, resulting in bone healing.

To further clarify the effect of exosomes on BMSC osteoblastogenesis *in vitro*, calcium deposition by N-BMSCs/DM-BMSCs treated with DM-Exos or N-Exos was investigated (Fig EV2 A). DM-Exos and N-Exos were derived from the BMSCs of DM and normal rats. The exosomes were cultured with BMSCs for 14 days, in each case. Calcium deposition was assessed using Alizarin red S staining. The results demonstrate that mineralization was significantly enhanced following treatment with both DM-Exos and N-Exos, although N-Exos treatment exhibited greater osteogenesis than DM-Exos treatment in both N-BMSC and DM-BMSC groups (Fig EV2 B,C). The results suggest that DM-Exos decreased osteoblastogenesis and calcium deposition in N-BMSCs while N-Exos increased osteoblastogenesis and calcium deposition in DM-BMSCs. N-Exos displayed a greater promotion of osteogenesis in DM-BMSCs than the treatment of N-BMSCs with DM-Exos.

### Exosomal miRNAs promote the osteogenesis of BMSCs

Given the negative impact of DM-Exos on bone formation and the positive effect of N-Exos on osteogenesis, the role of exosomal miRNA molecules in the osteogenesis of BMSCs was next evaluated. As is well known, exosomes are a type of extracellular vesicle containing miRNA molecules secreted from various cells. Circulating exosomes are considered an important mechanism for the transport of miRNA. To demonstrate that miRNAs are key functional components for osteogenesis to occur, a small interfering RNA (siRNA) molecule was used to knock down Drosha to inhibit miRNA synthesis and so deplete the secreted exosomes of miRNA molecules (Fig 4A). Firstly, N-BMSCs were treated with exosomes isolated from DM-BMSCs and N-BMSCs. Consistent with the previous results, the exosomes promoted Col-1, Runx-2, Sp7, and ALP expression, with greater osteogenesis caused by N-Exos than DM-Exos. In contrast, the treatment of N-BMSCs with exosomes isolated from DM-BMSCs and N-BMSCs in which Drosha had been knocked down demonstrates that these exosomes did not significantly promote osteogenesis to a greater extent than the control group. These results indicate that miRNAs are responsible for the promotion of osteogenesis by exosomes. DM-BMSCs were next treated with exosomes isolated from N-BMSCs and DM-BMSCs both with or without Drosha-knockdown. Consistent with the results above, exosomes without knockdown promoted Col-1, Runx-2, Sp7, and ALP expression in DM-BMSCs, although the N-Exos treatment group exhibited greater osteogenesis than the DM-Exos group. Conversely, the treatment of DM-BMSCs with exosomes isolated from N-BMSCs and DM-BMSCs that had undergone Drosha-knockdown indicated that the ability of the exosomes to promote osteogenesis in DM-BMSCs had disappeared (Fig 4B-F). These results indicate that miRNA molecules perform a critical role in the osteogenesis of exosomes.

**Fig 4.**
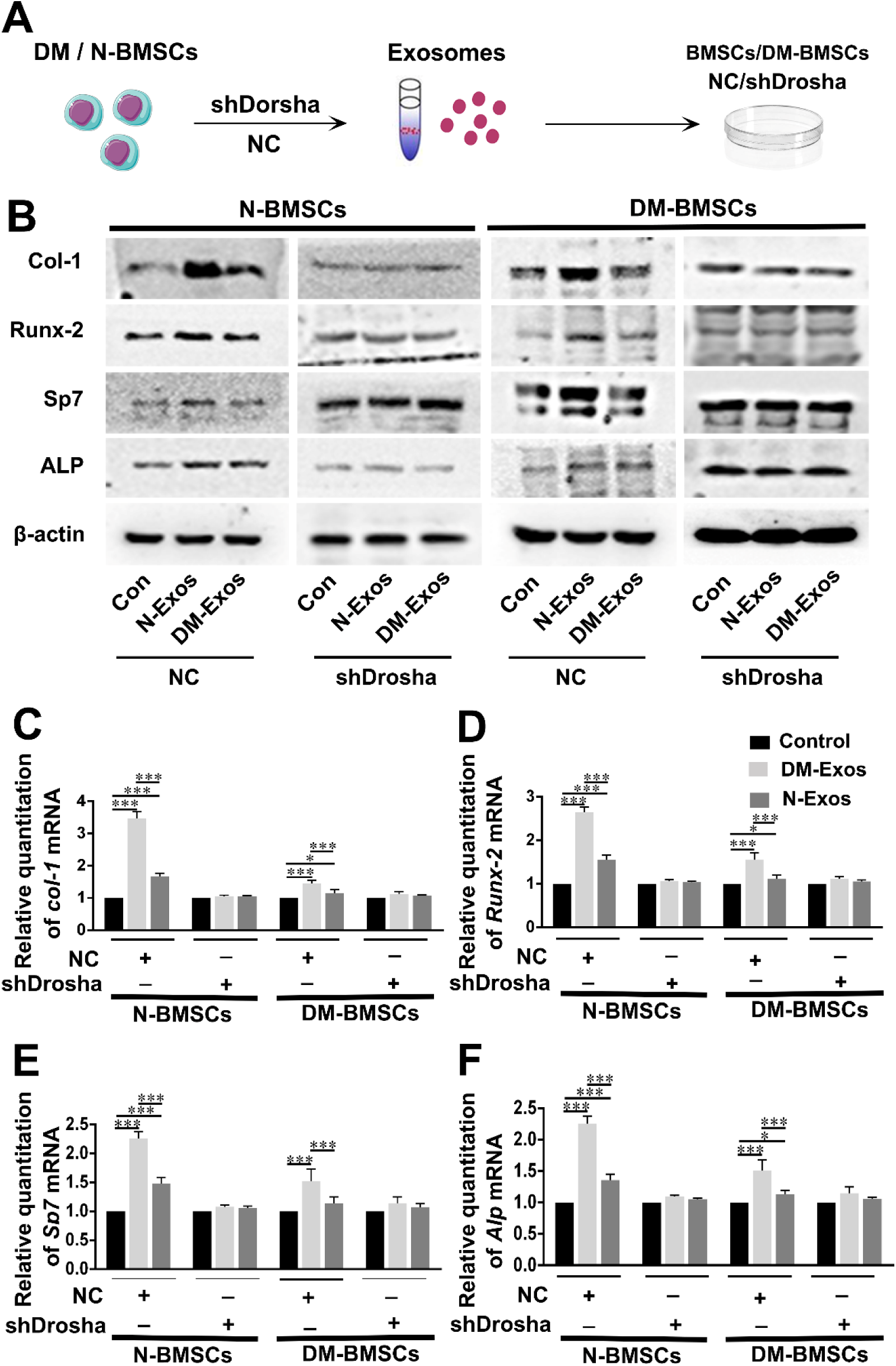
**Exosomes modulate BMSC differentiation through exosomal miRNAs** (A) Exos were collected from normal BMSCs and DM BMSCs after siRNA-mediated knockdown of Drosha and then cultured with untreated N-BMSCs or DM-BMSCs. After 48 h, the N-BMSCs and DM-BMSCs were collected and the protein expression levels of osteogenic molecules were measured by Western blotting. (B) Western blot analysis of Col-1, Runx-2, ALP, and Sp7 expression in N-BMSCs and DM-BMSCs cultured with Exos both with or without Drosha knockdown. n=3 in each group. (C-F) Quantitative analysis of Col-1, Runx-2, ALP, and Sp7 expression by Western blotting. n=3 in each group. *p < 0.05; **p < 0.01; ***p < 0.001. Data represent means ± SD.

Taken together, the results suggest that N-BMSCs secrete exosomal miRNAs in their normal state that provide protection against diabetes-induced bone loss.

### Hyperglycemia induces changes in BMSC-Exo miRNA expression

Given the negative osteogenic effects of exosomes from DM-BMSCs compared with those from N-BMSCs, DM-induced changes in the expression of miRNA molecules in BMSCs were then assessed. Deep sequencing of small RNAs from normal and DM BMSC-Exos was conducted to identify the differential expression of miRNAs. More than 600 miRNAs were identified in the exosomes of DM-BMSCs and N-BMSCs. Considerable differences in the miRNA expression of DM-Exos and N-Exos were observed. Of the differentially-expressed miRNAs, those most significantly differentially expressed in DM-Exos compared with N-Exos are presented in Fig 5A.

**Fig 5.**
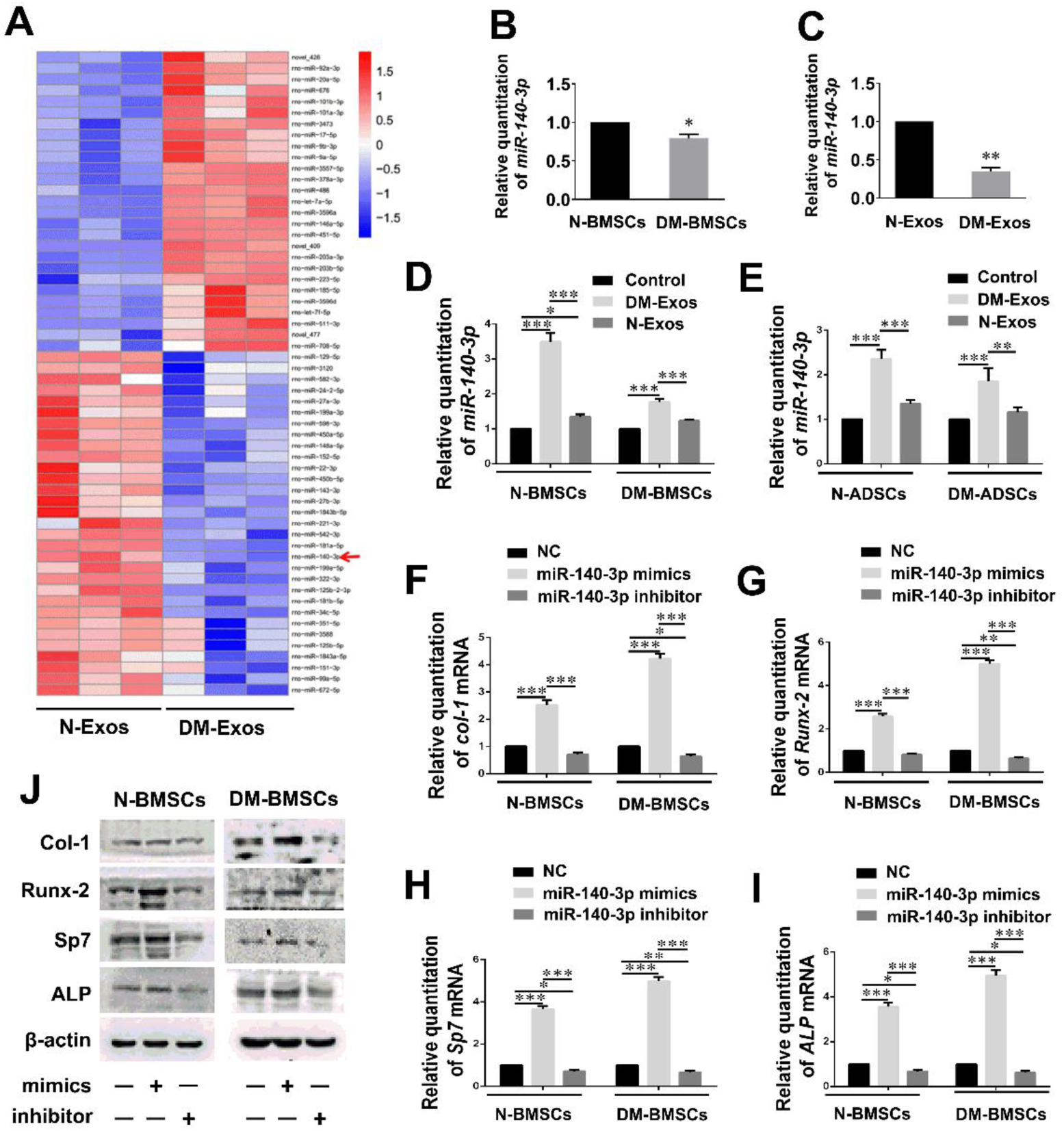
**Diabetes results in changes in the expression of miRNA in BMSCs-Exos** (A) Differential expression levels of exosomal miRNAs between N-Exos and DM-Exos. MiR-140-3p is an exosomal miRNA that underwent significantly reduced expression in DM-Exos compared with N-Exos. (B) Differential expression levels of miR-140-3p between N-BMSCs and DM-BMSCs. n=3 in each group. (C) Expression levels of miR-140-3p between N-Exos and DM-Exos. n=3 in each group. (D,E) MiR-140-3p expression in N-BMSCs/ADSCs and DM-BMSCs/ADSCs, which were treated with N-Exos and DM-Exos. n=3 in each group. (F-I) qPCR examination of mRNAs levels of *Col-1, Runx-2, ALP*, and *Sp7* in N-BMSCs and DM-BMSCs after miR-140-3p NC/mimics/inhibitor treatment. n=3 in each group. (J) Western blot analysis of Col-1, Runx-2, ALP and Sp7 expression in N-BMSCs and DM-BMSCs after miR-140-3p NC/mimics/inhibitor treatment. n=3 in each group. *p < 0.05; **p < 0.01; ***p < 0.001. Data represent means ± SD.

Among the miRNAs identified here, miR-140-3p, miRNA-34c-5p, miR-99a-5p, and miR-27a-3p were expressed in DM-Exos to a significantly greater extent than in N-Exos, and so were selected to evaluate the ability of miRNAs to influence osteogenesis in DM-BMSCs (Fig EV3). Mimics and inhibitors of the five miRNAs were evaluated to investigate their effect on osteoblastogenesis in DM-BMSCs. The results demonstrate that miR-140-3p significantly promoted osteogenesis in DM-BMSCs in contrast to the other miRNAs. Additionally, miR-140-3p has been previously reported to have anti-inflammatory effects(Min *et al*, 2015) and so the results suggest that miR-140-3p probably plays an important role in modulating the differentiation of BMSCs in DM.

MiR-140-3p expression in N-BMSCs was greater than in DM-BMSCs (Fig 5B), resulting in increased levels of miR-140-3p being present in N-Exos (Fig 5C). Incubation of N-BMSCs, DM-BMSCs, normal adipocyte-derived stem cells (N-ADSCs), and DM-ADSCs with N-Exos resulted in a significant increase in the expression of miR-140-3p within these cells, while treatment with DM-Exos led to a small or insignificant effect (Fig 5D,E). In addition, N-BMSCs were treated with miR-140-3p mimics/inhibitor to determine their effects on osteogenesis. qPCR analysis demonstrated that incubation of DM-BMSCs with miR-140-3p mimics resulted in Col-1, Runx-2, Sp7, and ALP expression in the miR-140-3p mimics treatment group being significantly higher than in the control and inhibitor groups. Furthermore, there was no significant difference between the control and inhibitor groups (Figs 5F-I). Western blot analysis yielded results consistent with the results described above (Fig 5J). MiR-140-3p mimics significantly promoted Col-1, Runx-2, Sp7, and ALP expression levels compared with those in the control and inhibitor groups. Additionally, no significant difference between the control and inhibitor groups was observed.

### MiR-140-3p promotes osteoblastogenesis in BMSCs via inhibition of the plexin B1/RhoA signaling pathway

To further explore the mechanism by which N-Exos induces osteogenesis, the influence of miR-140-3p on osteoblast differentiation signaling was determined. MiRNAs inhibit target gene functions by binding to the 3′ untranslated region (3′-UTR) or protein-coding sequence of specific mRNAs, assisting to degrade the mRNA or suppress translation. Bioinformatic software TargetScan was used to predict the possible downstream effectors of miR-140-3p (Fig EV4). Of the predicted target genes, Plxnb1 was selected as a candidate likely to influence bone formation. Plexin B1 has been reported to modulate osteogenesis by inhibiting the osteogenesis of osteoblasts and is activated by sema4D, which is secreted by osteoclasts. To examine the relationship between plxnb1 and miR-140-3p, miR-140-3p mimics and inhibitor were transfected into N-BMSCs and DM-BMSCs, respectively (Fig 6A). The results indicated that DM-BMSCs expressed a greater quantity of plexin B1 in contrast to N-BMSCs. However, when transfected with miR-140-3p mimics, the levels of plexin B1 expressed in DM-BMSCs and N-BMSCs decreased considerably. Consistent with Western blot analysis, qPCR demonstrated that miR-140-3p mimics substantially reduced plexin B1 expression (Fig 6B). To further ascertain the relationship and interaction between PLXNB1 and miR-140-3p, a dual Rluc/Fluc luciferase reporter plasmid containing the wildtype 3′-UTR of PLXNB1 (pSI-Check2-Plxnb1-3UTR) was generated (Fig 6C). HEK293T cells were transfected with pSI-Check2-Plxnb1-3UTR WT/MU dual Rluc/Fluc luciferase reporter plasmids and a miR-140-3p mimics/negative control. The results demonstrated that miR-140-3p significantly inhibited the luciferase signal caused by plxnb1 expression (Fig 6D). However, the inhibition was largely abolished when four crucial nucleotides in the putative binding site for miR-140-3p were mutated (Fig 6D). The results above suggest that plxnb1 may be the downstream target of miR-140-3p for the regulation of osteogenesis.

**Fig 6.**
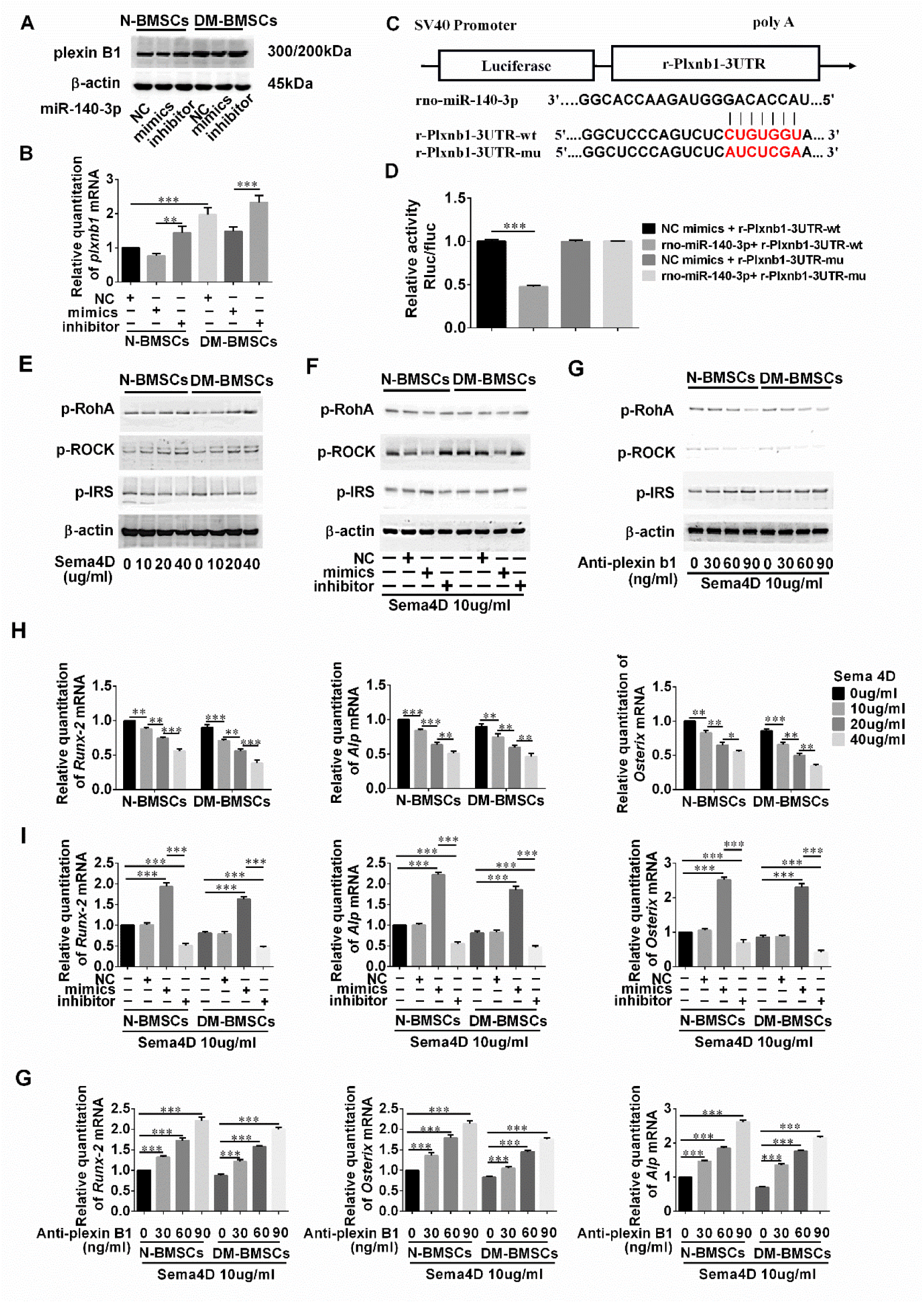
**MiR-140-3p regulation of BMSC differentiation via Plexin B1.** (A,B) Plexin B1 plays an important role in the interaction between osteoblasts and osteoclasts. The expression of plexin B1 in N-BMSCs and DM-BMSCs was examined using Western blot analysis and qPCR after treatment with miR-140-3p NC/mimics/inhibitor. n=3 in each group. (C) Schematic illustration of the sequences for miR-140-3p and the WT or mutated 3′-UTR of plxnb1 mRNA. (D) Dual Rluc/Fluc luciferase luminescence intensity of plxnb1 WT or mutated 3′-UTR reporter plasmids in HEK293 cells co-transfected with miR-140-3p mimics or miR-NC mimics. (E) Protein expression levels of p-RhoA, p-ROCK, and p-IRS in N-BMSCs and DM-BMSCs after Sema4D treatment (0, 10, 20, 40 μg/mL). (F) Levels of p-RhoA, p-ROCK, and p-IRS in N-BMSCs and DM-BMSCs transfected with miR-140-3p NC/mimics/inhibitor after treatment with Sema4D (10 μg/mL) as determined by Western blotting. (G) Levels of p-RhoA, p-ROCK, and p-IRS in N-BMSCs and DM-BMSCs as measured by Western blotting, after culture with anti-plexin B1 (0, 30, 60, 90 ng/mL) and Sema4D (10 μg/mL). (H) Levels of *Runx-2, ALP*, and *osterix(Sp7)* in N-BMSCs and DM-BMSCs after treatment with Sema4D (0, 10, 20, 40 μg/mL). (I) qPCR analysis of *Runx-2, ALP,* and *osterix (Sp7)* expression levels in N-BMSCs and DM-BMSCs transfected with miR-140-3p NC/mimics/inhibitor and cultured with Sema4D (10 μg/mL). (J) mRNAs expression levels of *Runx-2, ALP,* and *osterix (Sp7)* in N-BMSCs and DM-BMSCs treated with anti-plexin B1 (0, 30, 60, 90 ng/mL) and Sema4D (10 μg/mL). Data represent means ± SD (n=3 in each group. Western blotting and qPCR experiments were repeated 3 times). *p < 0.05; **p < 0.01; ***p < 0.001.

Semaphorin 4D (Sema4D) expressed by osteoclasts binds to its receptor Plexin-B1, resulting in the activation of the small GTPase RhoA, thereby inhibiting bone formation through inhibition of the insulin-like growth factor-1 (IGF-1) signaling pathway. Whether miR-140-3p regulates osteogenesis via plxnb1 or not was then further explored. Firstly, N-BMSCs and DM-BMSCs were incubated with sema4D, the agonist of plexin B1 (Fig 6E). The results demonstrate that the expression of p-RhoA and p-ROCK increased with increasing additions of sema4D, in a dose-dependent manner. In addition, p-IRS levels were reduced in these cells in a dose-dependent manner. Conversely, the expression of Runx-2, ALP, and Sp7 decreased due to increased sema4D in a dose-dependent manner in N/DM-BMSCs (Fig 6H). The results indicate that activation of the sema4D/plexin B1/RhoA/ROCK signaling pathway may inhibit osteoblastogenesis in BMSCs.

Secondly, the effects of miR-140-3p on the sema4D signaling pathway were investigated. MiR-140-3p mimics inhibited the effect of sema4D on the expression of p-RhoA and p-ROCK, and increased the expression of p-IRS in N/DM-BMSCs (Fig 6F). In addition, miR-140-3p mimics reversed the effects of sema4D on Runx-2, ALP, and Sp7 expression and enhanced their osteogenic effects (Fig 6I). These results suggest that miR-140-3p participates in osteoblast differentiation via modulation of the sema4D/plexin B1 signaling pathway. Lastly, BMSCs were incubated with anti-plexin B1 to elucidate the role of plexin B1 in osteogenesis. The results demonstrate that the expression of p-RhoA and p-ROCK decreased with increasing levels of anti-plexin B1 in a dose-dependent manner, despite treatment with sema4D (Fig 6G). In addition, the expression of Runx-2, ALP, and Sp7 increased relative to the dose of anti-plexin B1 in N/DM-BMSCs (Fig 6J). Thus, the results suggest that plxnb1 is the direct downstream mRNA target of miR-140-3p, and that miR-140-3p can rescue osteogenesis by inhibition of the sema4D/plexin B1 signaling pathway in DM-BMSCs.

### Exosomes derived from BMSCs overexpressing miR-140-3p promote osteogenesis that heals bone defects when experiencing hyperglycemia

The therapeutic effects of miR-140-3p in regulating bone formation in DM rats were investigated by producing BMSC-specific exosomes overexpressing miR-140-3p and transplanting them into the bone defects of DM rats. To obtain exosomes overexpressing miR-140-3p, BMSCs were transfected with a lentivirus that overexpressed miR-140-3p (Fig EV5). Exosomes overexpressing miR-140-3p were collected by ultracentrifugation. qPCR was used to confirm that, compared with N-Exos and DM-Exos, exosomes from BMSCs transfected with the lentivirus displayed high levels of miR-140-3p, with osteogenic properties of the BMSCs consistent with those levels (Fig EV5).

DM rats with bone defects were transplanted with 240 μg N-Exos, DM-Exos, or exosomes overexpressing miR-140-3p (140-Exos) (Fig 8A). Micro-CT was used to quantify bone mass in the different groups (Fig 7A,B). The results demonstrate that although each exosomal treatment increased bone mass, the rats in which 140-Exos were transplanted experienced an enhancement in bone mass greater than that observed in the N/DM-Exos transplanted rats after both 2 and 8 weeks, as manifested in the BV/TV values. The BV/TV values after 8 weeks were higher than after 2 weeks in the 140-Exos transplantation group. Trabecular number and trabecular thickness increased after all exosome treatments compared with the control group, although their values were greater in the 140-Exos treatment compared with those for the N and DM-Exos treatments. Similarly, trabecular number and thickness were also higher after 8 weeks compared with after 2 weeks in the 140-Exos treated rats. Trabecular spacing declined in the exosome treatment groups compared with values in the control group. The data described above indicate that 140-Exos promoted bone defect remodeling which demonstrated the osteogenic effect of miR-140-3p.

**Fig 7.**
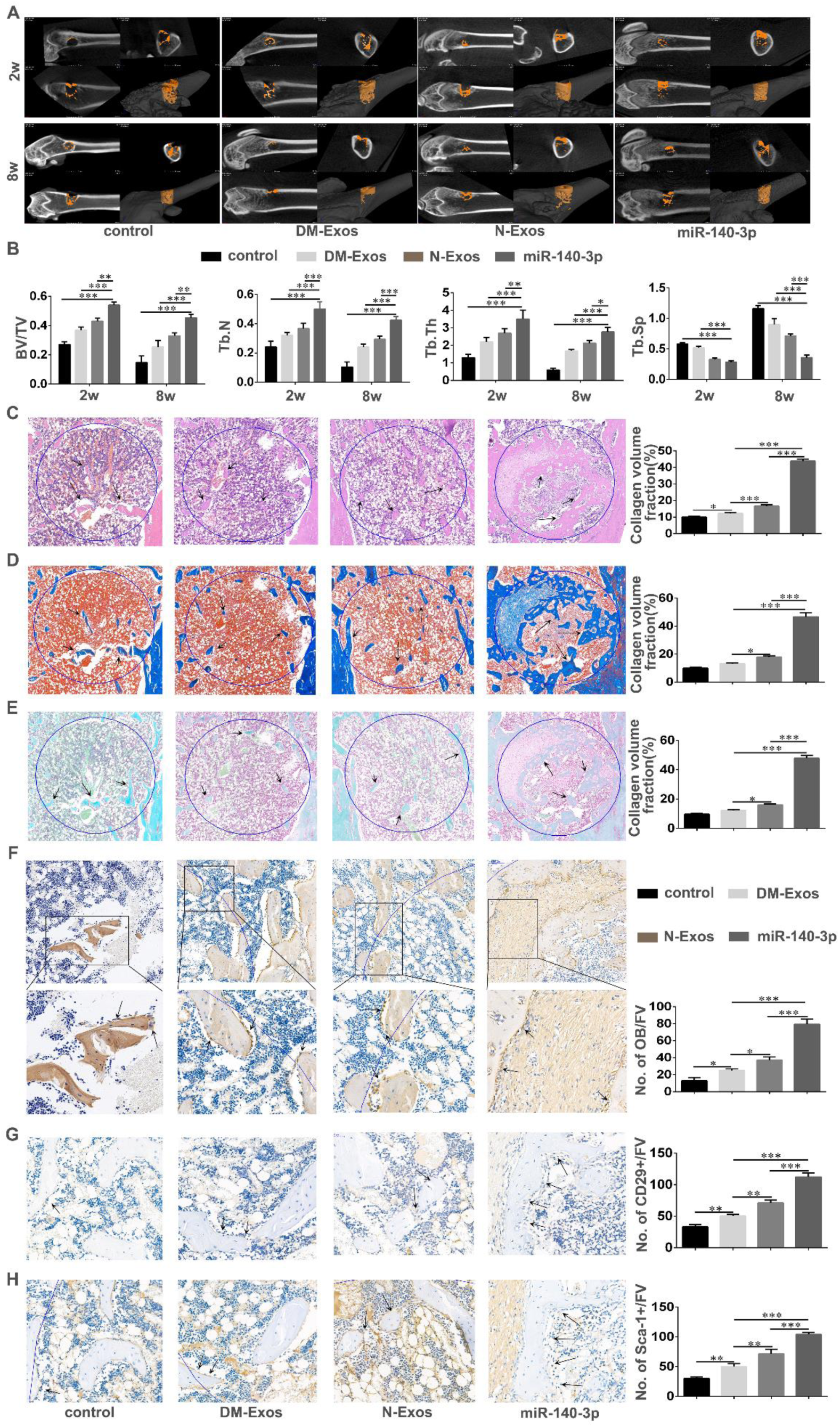
**MiR-140-3p promotes bone formation in DM rats with bone defects** (A) Representative micro-CT images within a region of interest (ROI) comprising the bone defect after 2 and 8 weeks following surgery in the bone defect of normal rats and those with diabetes mellitus after treatment with N-Exos, DM-Exos, and Exos overexpressing miR-140-3p. n=5 in each group. (B) Quantitative analysis of new bone formation from micro-CT imaging in DM rats with bone defects after 2 and 8 weeks after transplantation with N-Exos, DM-Exos, or Exos overexpressing miR-140-3p. BV: bone volume; TV: tissue volume; Tb,N: trabecular number; Tb.Th: trabecular thickness; Tb.Sp: trabecular Spacing. n=5 in each group. (C-E) Histological analysis of decalcified sections of bone defects in diabetic rats treated with N-Exos, DM-Exos, or Exos overexpressing miR-140-3p. HE/Masson’s/Safranin O/fast green staining of the new bone formation within the defect. Black arrows indicate new bone stained in light pink/blue/green within the 3mm-diameter defect marked as a circle. Quantitative analysis of new bone formation was performed using ImageJ software. n=5 in each group. (F-H) Immunohistological analysis of osteoblasts and BMSCs in diabetic rats treated with N-Exos, DM-Exos, or Exos overexpressing miR-140-3p. Collagen-1 and CD29/Sca-1 staining of osteoblasts and BMSCs in sections containing the bone defect. Black arrows indicate osteoblasts and BMSCs on the bone surface. n=5 in each group; *p < 0.05; **p < 0.01; ***p < 0.001. Data represent means ± SD.

In hematoxylin and eosin-stained sections, the extent of light pink collagen in the bone defect rats with Exos treated was greater than in control rats. Consistent with the findings above, the volume of collagen in the 140-Exos rats was highest in the N-Exos and DM-Exos transplantation groups (Fig 7C). In the Masson’s-stained sections, blue collagen staining was observed. The volume of mineralized bone tissue in the 140-Exos transplantation group was significantly greater than in the N/DM-Exos groups. In addition, the volume of mineralized bone tissue in the control rats was less than that observed in the exosome-treated rats (Fig 7D). Safranin O/Fast green staining demonstrated that the volume of mineralized bone in 140-Exos was larger than that observed in the N/DM-Exos groups (Fig 7E). Immunohistochemical staining of collagen-1 indicated that the numbers of osteoblasts in the control group were the least of all four groups while the numbers of osteoblasts in the 140-Exos group were greatest of the N/DM/140-Exos groups (Fig 7F). The expression of the BMSC markers CD29/Sca-1 indicated that there were fewer BMSCs on the bone surface in the N/DM-Exos treatment groups than in the 140-Exos treated group, but more than in the control group (Fig 7G,H). The data above indicate that 140-Exos significantly accelerated bone formation by increasing the numbers of BMSCs and osteoblasts in DM rats, resulting in greater mineralized bone formation and bone defect healing.

## Discussion

Delayed diabetic bone healing already represents an intractable medical problem, as current strategies are functionally limited. In the present study, a series of *in vitro* and *in vivo* studies were performed to determine whether BMSC-derived exosomal miR-140-3p promotes bone formation, and thus confirm the paradigm in which miRNA regulates the homeostatic mechanism of the local bone environment.

We found that DM-Exos and N-Exos derived from BMSCs containing miRNA modulated bone formation both *in vivo* and *in vitro*. In *in vitro* studies, exosomes from N-BMSCs and DM-BMSCs were found to promote osteoblastogenesis and mineralization in the BMSCs from both normal and DM rats, although N-Exos had a greater osteogenesis effect on osteoblastogenesis than did DM-Exos. Similarly, both N-Exos and DM-Exos enhanced bone regeneration in a femoral defect model in normal and DM rats, although the bone regenerative effect of N-Exos was greater than DM-Exos *in vivo*.

Bone modeling and remodeling describe processes in which bones are shaped or reshaped by the independent action of osteoblasts and osteoclasts. Osteoclasts resorb bone and release osteogenic factors into the bone marrow. In addition, BMSCs not only function as progenitors of osteoblasts but also support osteoclast differentiation (Xu *et al*, 2012). Diabetes has been shown to alter the properties of bone and impair bone repair in both humans and animals. There is accumulating evidence that diabetes mellitus decreases bone formation and the osteogenic capability of BMSCs (Jiao *et al*., 2015). DM is a chronic metabolic disease leading to high blood glucose levels which contribute to the production of advanced glycation end products (AGEs), reactive oxygen species, and tumor necrosis factor (Compston, 2018; Cruz *et al*, 2013; Donath *et al*, 2019). AGEs bind to specific receptors on BMSCs and inhibit BMSC differentiation, proliferation, and migration (Notsu *et al*, 2014). Additionally, AGEs increase BMSC apoptosis via MAPK signaling and oxidative stress (Kume *et al*, 2005; Yang *et al*, 2010). With high levels of inflammatory factors and the inhibitory action of hyperglycemia, the capacity of BMSCs to form bone has been shown to become downregulated with increased apoptosis, leading to a disruption of bone coupling and remodeling (Alblowi *et al*, 2009; Claes *et al*, 2012).

Exosomes are derived from a variety of cell types and transfer into target cells whereby various functions are activated. Increasing numbers of reports indicate that stem cell-derived exosomes have the capability to promote bone formation (Liu *et al*, 2017a). As exosomes from different cell types provide a variety of therapeutic effects due to their specific cargos, we believe that exosomal therapy has considerable potential for clinical translation (Chen *et al*, 2017; Hao *et al*, 2017). In the present study, exosomes were extracted from BMSC-conditioned media by ultracentrifugation and identified from other extracellular vesicles by their characteristic diameter of 30-100 nanometers, as measured by NanoSight nanoparticle analysis and morphology by electron microscopy. Additionally, the exosome-specific marker TSG101 and extracellular vesicle-related protein markers HSP70, CD63, and CD9 were measured by Western blot analysis. Together, the results indicate that the extracellular vesicles were indeed exosomes (Peinado *et al*, 2012; Su *et al*, 2019; Ying *et al*, 2017). Moreover, we used transwell culture and transfection to demonstrate that exosomal miRNAs could directly transfer into recipient cells. We firstly transfected BMSCs with Cy3-labeled miR-223 and transplanted them into the upper chamber of a transwell insert with untransfected BMSCs placed in the lower chamber. The red fluorescence of Cy3-labeled miR-223 in the BMSCs in the lower chamber was detected. Untransfected BMSCs were then incubated with exosomes labeled with PKH26 fluorescent dye. This demonstrated that non-transfected BMSCs absorbed fluorescent-labeled exosomes, displaying green fluorescence. miR-223 expression also increased in non-transfected BMSCs, which was inhibited by prior treatment of BMSCs with GW4869 (an inhibitor of EV secretion). The results demonstrated that exosomes and exosomal miRNA could be directly transferred into recipient cells.

The bone environment has also been found to be altered in individuals suffering from T2DM, which has a negative impact on bone formation. We found that DM-BMSC derived-Exos promoted bone formation and increased bone mass in normal rats compared with control rats, although the osteogenic capability of DM-Exos was significantly impaired compared with that of N-Exos treated rats. Nevertheless, N-Exos promoted bone regeneration and increased bone volume/tissue volume in DM rats compared with DM-Exos transplanted rats. Similarly, the volume of collagen in DM-Exos-treated normal rats was less than that observed in the N-Exos treatment group, while DM-Exos decreased the number of osteoblasts and BMSCs in normal rats compared with the N-Exos group. As described above, N-Exos enhanced the volume of collagen in DM rats and the numbers of osteoblasts and BMSCs compared with the DM-Exos group. In addition, exosomes promoted mineralization generated in BMSCs, although the osteogenic capability of N-Exos was superior to that of DM-Exos. The results indicate that DM-BMSC-derived exosomes impaired bone regeneration. The results were consistent with previous studies (Zhu *et al*., 2019). However, few studies have concentrated on differences between DM-BMSC-derived exosomes and those from normal BMSCs. Thus, we explored the differences here.

Exosomal miRNAs have become a focus of recent research attention due to their numerous roles in regulation (Ying *et al*, 2021; Ying *et al*., 2017). MiRNAs and RNA-induced silencing complexes (RISCs) are packaged into exosomes and transferred to recipient cells to silence mRNAs (Valadi *et al*, 2007). Numerous research studies have revealed that a number of bone-derived exosomal miRNAs participate in bone remodeling. There are 43 exosomal miRNAs highly expressed in MC3T3-E1 cells during the osteogenesis process, such as miR-30d-5p and miR-133b-3p (Li *et al*, 2008; Zhang *et al*, 2011). Moreover, miR-135b, miR-203, and miR-299-5p are upregulated in human-BMSCs derived exosomes during osteogenesis (Panganiban *et al*, 2016; Salazar-Roa *et al*, 2020; Schaap-Oziemlak *et al*, 2010). Thus, in the present study, we aimed to identify the differences in exosomal miRNAs between N-Exos and DM-Exos. To investigate the effect of DM on changes to miRNAs, high-throughput sequencing was used to determine the identity of differentially expressed exosomal miRNAs. More than 600 miRNAs were sequenced and quantified, from which miR-140-3p, miRNA-34c-5p, miR-99a-5p, and miR-27a-3p were found to be in significantly different abundance in DM-Exos compared with N-Exos. These have been identified in previous studies for their ability to promote osteogenesis (Fu *et al*, 2019; Gandla *et al*, 2017; Xie *et al*, 2017). BMSCs were incubated with miRNA mimics and inhibitor to ascertain the osteogenic effects of these miRNAs. The results indicated that miR-140-3p displayed the greatest promotion of BMSC differentiation compared with other miRNAs. In addition, the expression of miR-140-3p was lower in DM-BMSC, DM-ADSC, and DM-BMSC Exos, suggesting that the inhibition by DM of osteogenesis in BMSCs may decrease levels of miRNA in BMSCs. To further demonstrate the impact of miR140-3p on osteogenesis, we transfected BMSCs with miR-140-3p, the results of which demonstrated that miR-140-3p mimics significantly promoted Col-1, Runx-2, Sp7, and ALP expression in DM-BMSCs and N-BMSCs compared with the control and inhibitor groups. Western blot analysis and qPCR indicate that miR-140-3p played an important role in the differentiation of BMSCs. Similarly, Rakefet Pando found that chondrocyte-specific miR-140-3p displayed the greatest expression in mature EGP(epiphyseal growth plate), and was one of only a few miRNAs that were significantly reduced in expression following restricted nutrition. Additionally, SIRT1 levels increased significantly in nutrition-restricted states. Their study indicated that the expression of miR-140-3p was positive during nutrition-induced catch-up growth in the epiphyseal growth plate (EGP) (Pando *et al*, 2012). Moreover, consistent with the findings of the present study, S. Takahara found that miR-140-3p levels were significantly lower in DM rats after 14 days compared with control rats (Takahara *et al*, 2018). Furthermore, miR-140-3p was found to be significantly highly expressed in rats undergoing bone fracture healing compared with those that were not healing (Waki *et al*, 2016). Together, the results established that miR-140-3p exerted an osteogenic effect on BMSCs.

To further determine the mechanisms by which miRNA-140-3p was involved in the regulation of bone regeneration, we explored the target mRNAs of miR-140-3p. Theoretically, miRNAs can have more than one mRNA target, and there could be more than one miRNA directed against an mRNA molecule (Bartel, 2009). We used TargetScan software to analyze the sequenced mRNA molecules in BMSCs. This analysis revealed that several mRNA molecules represented downstream effectors of miR-140-3p. Of the numerous candidates, plxnb1 emerged as having a positive association with bone formation and osteogenesis. Plexin B1 has been reported to be a receptor of sema4D, which is secreted by osteoclasts. Sema4D binds to plexin B1 and inhibits osteogenesis in osteoblasts, reducing bone formation (Deb Roy *et al*, 2017; Negishi-Koga *et al*, 2011; Terpos *et al*, 2018). To confirm that plxnb1 is the effector of miR-140-3p, dual Rluc/Fluc luciferase reporter plasmids containing the wildtype 3′-UTR of PLXNB1 (pSI-Check2-Plxnb1-3UTR) were generated, the results showing that miR-140-3p significantly inhibited luciferase activity due to plxnb1. However, this inhibition was largely abolished when four crucial nucleotides were mutated in the putative binding site of miR-140-3p. In addition, the expression of plexin B1 was significantly lower in BMSCs transfected with miR-140-3p. The results demonstrate that miR-140-3p promoted osteogenesis by inhibition of the plexinB1/RhoA/ROCK signaling pathway.

In summary, the findings indicate that considerable differences exist in the miRNAs contained in N-Exos compared with DM-Exos, which supports the hypothesis that BMSC-derived exosomal miR-140-3p functions as an important regulator of osteoblastogenesis. Moreover, administration of miR-140-3p to the bone marrow microenvironment may provide a potential therapeutic strategy for diabetes-induced bone fracture/defect.

## Materials and Methods

### Animals and Treatment

One hundred 4-week-old male Sprague-Dawley rats were obtained from the Animal Center of the Air Force Military Medical University (Xi’an, China). All animal experiments and care protocols were reviewed and approved by the Animal Research Committee of the Air Force Military Medical University ( SCXK-2019-001 ). All animals were provided *ad libitum* access to food and water prior to the experiments. Rats were fed a high fat and high carbohydrate diet (HFD) for one month after which streptozotocin (Sigma-Aldrich, St. Louis, MO, USA, 40 mg/kg) dissolved in 10 mmol/L citrate buffer (pH 4.5) was injected intraperitonealy once into the rats, which were then fasted for 2 h, then allowed to eat freely. After 7 days, blood glucose concentrations were tested via a blood glucometer. Rats with blood glucose levels higher than 16.7 mmol/L were considered to be diabetes mellitus positive.

### Bone defect model

A bone defect model was established in accordance with a previous study. Nine-week-old male Sprague-Dawley rats were anesthetized using sodium pentobarbitone (3% m/m). A 1.0-cm-long incision was created along the femur in the sagittal plane on right side. A bone defect was created in the metaphysis by drilling a 3 mm-diameter hole. Exosomes (240 μg) were thoroughly mixed with Matrigel (Corning, NY, USA, 356230) and placed on ice prior to injecting into the bone defects. Forty rats were randomly allocated into the following four groups: (a) matrigel group (control group; n=10); (b) matrigel mixed with N-Exos (N-Exos group; n=10); (c) matrigel mixed with DM-Exos (DM-Exos group; n=10); and (d) matrigel mixed with Exos overexpressing miR-140-3p (140-Exos group; n=10).

### Micro-CT Analysis

Rats were anesthetized with 3% sodium pentobarbitone allowing the microarchitecture of the femoral cavities to be examined by micro-CT. Bone images were reconstructed using an isotropic voxel size of 10 μm. Bone volume/total volume, trabecular thickness, trabecular number, and trabecular spacing values were recorded to evaluate bone formation. Three-dimensional imaging and the reconstruction of the microarchitecture were performed using an Inveon Research Workplace 2.2 (SIEMENS Healthineers, Berlin, Germany).

### BMSC Isolation and Treatment

BMSCs were harvested from both femurs of the rats and sorted using a FACS Aria II flow cytometer (BD Biosciences, New Jersey, USA). Bone marrow cells were flushed from the bone marrow cavity after which a red blood cell lysis buffer was used to eliminate red blood cells. The remaining cells were then enumerated. Positivity toward a PE-conjugated mouse monoclonal antibody against integrin beta 1 (Abcam, Cambridge, UK, ab218273) and a FITC-conjugated rat monoclonal antibody against Sca1/Ly6A/E (Abcam, ab268016) was utilized to separate BMSCs from other bone marrow cells. The sorted cells were transfected with miRNA mimics/inhibitor, or treated with siRNA or an agonist for additional experiments. In addition, exosomes (2 μg) were added to the culture medium of BMSCs (1×10^5^).

### Exosomes Isolation, Identification

Exosomes were isolated using ultracentrifugation, as described previously(Su *et al*., 2019). Cell culture supernatants were centrifugated at 300 g for 10 min, 2000 g for 30 min, 10,000 g for 5 min at 4 ℃, then finally at 100,000 g for 4 h at 4 ℃ using an SW28 rotor (Beckman Coulter, Fullerton, CA, USA). The pelleted exosomes were collected then subsequently resuspended in PBS. Western blotting was used to quantify specific marker proteins in the exosomes, including HSP70, TSG101, CD63, and CD9. The size distribution of the exosomes was measured by nanoparticle tracking analysis (ZetaView PMX 110 ,Meerbusch, Germany) and their morphology was assessed using transmission electron microscopy (FEI Tecnai Spirit, Oregon, USA).

### Immunohistochemistry

Harvested rat femurs were fixed in 4% paraformaldehyde (PFA) then decalcified in ethylenediaminetetraacetic acid (EDTA; 10%, pH 7.0) prior to embedding in paraffin. Samples were sliced into 5 μm-thick sections using a microtome in a plane parallel to that of the bone defect cavity. The sections were then incubated in dimethylbenzene I and II, twice in 100% ethanol, twice in 95% ethanol, then in 90% ethanol and 80% ethanol, respectively. Endogenous peroxidase activity was eliminated with 3% H_2_O_2_ after which samples were incubated with goat serum at room temperature for 30 min. Finally, the samples were incubated with a specific primary antibody for 12 h at 4°C, from the following panel: mouse monoclonal anti-collagen I (Col-1;ab270993, Abcam); rabbit polyclonal anti-CD29 (GeneTex,California,USA,GTX128839); rabbit anti-mouse Sca-1 (Biolegend, California,USA,108103). Sections were then incubated with a biotinylated secondary antibody or conjugated with horseradish peroxidase (HRP) at room temperature for 30 min, after which color was developed with diaminobenzidine (DAB) and the sections counterstained with hematoxylin. Images were acquired following observation by light microscopy (Olympus BX53, Tokyo, Japan).

### Histological Analysis

The harvested rat femurs were fixed in PFA (4%) then decalcified in EDTA (10%, pH 7.0), respectively, prior to embedding in paraffin. The tissue was sliced into 5 μm-thick sections using a microtome in a plane parallel to that of the bone. The sections were deparaffinized then rehydrated, as reported previously(Wang *et al*, 2020). The sections were then stained with hematoxylin and eosin, Masson’s trichrome, and Safranin O/fast green to evaluate new bone formation.

### Alizarin staining

Alizarin red S staining was used to evaluate mineralization. The BMSCs were fixed in 4% paraformaldehyde for 30 min, then stained with alizarin red S for 20 min. Images were obtained via an Olympus BX53 microscope (Olympus, Tokyo, Japan). The extent of calcium deposition was assessed in each group to determine osteogenesis.

### Western blotting

Cells or exosomes were lysed in RIPA buffer and the concentration of protein in the lysates was measured using a bicinchoninic acid (BCA) kit. The lysates were diluted with loading buffer (5×) then heated to 100℃ for 10 min. Individual proteins were separated using 10% sodium dodecyl sulfate polyacrylamide gel electrophoresis (SDS-PAGE) after which they were transferred onto polyvinylidene fluoride (PVDF) membranes PVDF, Millipore, MA, USA). Non-specific binding was blocked by incubating the membranes with 5% nonfat milk for 2 h at room temperature and the identity of proteins evaluated by incubation at 4 ℃ overnight with the following primary antibodies (Abcam): HSP70 (1:1000, ab2787), TSG101 (1:5000, ab125011), CD63 (1 μg, ab193349), CD9 (1:2000, ab92726), col-1 (1:1000, ab270993), Runx-2 (1:1000, ab76956), Sp7 (1:1000, ab209484), and ALP (1:1000, ab229126). Membranes were then incubated with either an anti-rabbit IgG or anti-mouse IgG (1:10000, Boster Bio-Technology, Wuhan, China) HRP-conjugated secondary antibody, as appropriate, for 2 h at room temperature. The density of the immunoreactive bands was analyzed using a ChemiDoc XRS (Bio-Rad, Hercules, CA, USA) and quantified using Quantity One version 4.1.0 software (Bio-Rad).

### qPCR

Total RNA from exosomes or BMSCs was extracted using Trizol reagent ((Invitrogen, California, USA)). A One Step SYBR® PrimeScript™ qPCR kit (TaKaRa Bio, Otsu, Japan) was used to synthesize cDNA, in accordance with the manufacturer’s instructions. Quantitative real-time PCR (qPCR) was performed using SYBR® Premix Ex Taq™ (TaKaRa) in a Bio-Rad CFX96™ real-time PCR system using the following thermocycling conditions: pre-denaturation at 95 °C for 5 s, followed by 40 cycles of denaturation at 95 °C, 10 s; annealing at 57 °C, 20 s; and extension at 72 °C, 20 s. The relative expression of the specified genes was calculated using the 2^−ΔΔCT^ method after normalization to GAPDH expression.

In addition, miRNA was quantified by synthesizing cDNA using a Sangon Biotech miRNA First Strand cDNA synthesis (tailing reaction) kit (B532451), in accordance with the manufacturer’s protocol then performing qPCR using a Sangon Biotech microRNA qPCR kit with SYBR Green (B532461) in a Bio-Rad CFX96™ real-time PCR detection system (Bio-Rad) using the following thermocycling conditions: pre-denaturation at 95 °C for 30 s, followed by 40 cycles of denaturation at 95 °C, 5 s; annealing at 60 °C, 30 s; and extension at 72 °C, 30 s. The relative expression of each gene was calculated using the 2^−ΔΔCT^ method after normalization to U6 expression.

### Firefly luciferase & Renilla luciferase activity assay

HEK293 cells were seeded in 96 well plates an cultured to 50%-70% confluence prior to transfection. A 0.16 μg quantity of plasmids consisting of the 3′-untranslated region (UTR) of plxnb1 with a firefly luciferase reporter and psiCHECK-2 with Renilla luciferase reporter were mixed with 10 μL DMEM and 5pmol rno-miR-140-3p/Negative Control (N.C.) at room temperature for 5 min, termed solution A, then 10 μL DMEM were mixed with 0.3 μL Lipofectamine MessengerMax (ThermoFisher, Massachusetts, USA, LMRNA001), termed solution B. Solution C was formed by blending A with B and incubating at room temperature for 20 min. The HEK293 cells were then incubated with solution C for 6 h after which luciferase luminescent intensity was measured using a Promega dual-luciferase assay kit, in accordance with the manufacturer’s protocol.

### Statistical analysis

All data are presented as means ± standard deviation (SD). Three independent experiments were analyzed using SPSS version 15.0 software. Statistical significance was determined by one-way ANOVA, while multiple comparisons were performed using a Student-Newman-Keuls t-test. P < 0.05 was considered statistically significant.

## Acknowledgements

This work was supported by the National Key Research and Development Program of China (grant number 2017YFC1104900), the National Natural Science Foundation of China (grant numbers 51871239, 51771227 and 81772328), the Shaanxi Key Research and Development Program (grant number 2020SF-088) and China Postdoctoral Science Foundation (2020M673663)

## Author Contributions

Conceptualization, Ning Wang and Zheng Guo; Methodology, Ning Wang, Xue Ma, and Xiaokang Li; Investigation, Xuanchen Liu, Xinghui Wei, Zhen Tang and Yichao Liu; Writing – Original Draft, Ning Wang, Hui Dong, Hao Wu and Zhigang Wu; Writing –Review & Editing, Xue Ma and Xiaokang Li; Funding Acquisition, Zheng Guo, Xiaokang Li and Xue Ma; Resources,Ning Wang and Zheng Guo; Supervision, Zheng Guo, Xue Ma, Xiaokang Li.

## The Paper Explained Problem

Impaired bone healing is an important complication associated with diabetes mellitus, which affecting millions of people around the world. Aging, gender and the dysfunction of hormone contribute to unsatisfying bone restoration in diabetes mellitus. Hyperglycemia decreases BMSCs proliferation and differentiation, but the precise mechanisms are not yet fully understood.

## Results

We found that exosomes secreted by BMSCs derived from type 2 diabetic rats (DM-Exos) delayed the healing of bone defects in normal rats while those from normal rats (N-Exos) promoted bone formation in diabetic rats. High-throughput sequencing method was conducted to identify the differential expression of miRNAs between DM-Exos and N-Exos and miR-140-3p was identified as a significant miRNA in modulating BMSCs differentiation in diabetes mellitus for promoting Col-1, Runx-2, Sp7, and ALP expression. Bioinformatic software TargetScan predicted *Plxnb1* was the downstream effectors of miR-140-3p. Dual Rluc/Fluc luciferase reporter showed that miR-140-3p significantly inhibited luciferase activity due to *plxnb1*. However, this inhibition was largely abolished when four crucial nucleotides were mutated in the putative binding site of miR-140-3p. In addition, Sema4D (agonist of plexin B1) could significantly decrease the expression of Runx-2, Osterix and ALP expression in BMSCs with miR-140-3p overexpression. Besides, we found that miR-140-3p promoted osteogenesis by inhibition of the plexinB1/RhoA/ROCK signaling pathway. Furthermore, rats transplanted with exosomes overexpression miR-140-3p were experienced an enhancement in bone mass greater than that observed in the N/DM-Exos transplanted rats.

## Impact

This study indicates that considerable differences exist in the miRNAs contained in N-Exos compared with DM-Exos, which supports the hypothesis that BMSC-derived exosomal miR-140-3p functions as an important regulator of osteoblastogenesis in diabetes mellitus. Moreover, administration of miR-140-3p to the bone marrow microenvironment may provide a potential therapeutic strategy for diabetes-induced bone fracture/defect.

## Data availability

This study includes no data deposited in external repositories. The data that support the findings of this study are available from the corresponding author upon reasonable request.

## References

Alblowi J, Kayal RA, Siqueira M, McKenzie E, Krothapalli N, McLean J, Conn J, Nikolajczyk B, Einhorn TA, Gerstenfeld L et al (2009) High levels of tumor necrosis factor-alpha contribute to accelerated loss of cartilage in diabetic fracture healing. Am J Pathol 175: 1574–1585

Ameres SL, Martinez J, Schroeder R (2007) Molecular basis for target RNA recognition and cleavage by human RISC. Cell 130: 101–112

Bartel DP (2004) MicroRNAs: genomics, biogenesis, mechanism, and function. Cell 116: 281–297

Bartel DP (2009) MicroRNAs: target recognition and regulatory functions. Cell 136: 215–233

Bjorge IM, Kim SY, Mano JF, Kalionis B, Chrzanowski W (2017) Extracellular vesicles, exosomes and shedding vesicles in regenerative medicine - a new paradigm for tissue repair. Biomater Sci 6: 60–78

Brennecke J, Stark A, Russell RB, Cohen SM (2005) Principles of microRNA-target recognition. PLoS Biol 3: e85

Chen B, Li Q, Zhao B, Wang Y (2017) Stem Cell-Derived Extracellular Vesicles as a Novel Potential Therapeutic Tool for Tissue Repair. Stem Cells Transl Med 6: 1753–1758

Claes L, Recknagel S, Ignatius A (2012) Fracture healing under healthy and inflammatory conditions. Nat Rev Rheumatol 8: 133–143

Compston J (2018) Type 2 diabetes mellitus and bone. J Intern Med 283: 140–153

Cruz NG, Sousa LP, Sousa MO, Pietrani NT, Fernandes AP, Gomes KB (2013) The linkage between inflammation and Type 2 diabetes mellitus. Diabetes Res Clin Pract 99: 85–92

Deb Roy A, Yin T, Choudhary S, Rodionov V, Pilbeam CC, Wu YI (2017) Optogenetic activation of Plexin-B1 reveals contact repulsion between osteoclasts and osteoblasts. Nat Commun 8: 15831

Donath MY, Dinarello CA, Mandrup-Poulsen T (2019) Targeting innate immune mediators in type 1 and type 2 diabetes. Nat Rev Immunol 19: 734–746

Fu YC, Zhao SR, Zhu BH, Guo SS, Wang XX (2019) MiRNA-27a-3p promotes osteogenic differentiation of human mesenchymal stem cells through targeting ATF3. Eur Rev Med Pharmacol Sci 23: 73–80

Gandla J, Lomada SK, Lu J, Kuner R, Bali KK (2017) miR-34c-5p functions as pronociceptive microRNA in cancer pain by targeting Cav2.3 containing calcium channels. Pain 158: 1765–1779

Garcia-Garcia A, de Castillejo CL, Mendez-Ferrer S (2015) BMSCs and hematopoiesis. Immunol Lett 168: 129–135

Gomez-Barrena E, Rosset P, Gebhard F, Hernigou P, Baldini N, Rouard H, Sensebe L, Gonzalo-Daganzo RM, Giordano R, Padilla-Eguiluz N et al (2019) Feasibility and safety of treating non-unions in tibia, femur and humerus with autologous, expanded, bone marrow-derived mesenchymal stromal cells associated with biphasic calcium phosphate biomaterials in a multicentric, non-comparative trial. Biomaterials 196: 100–108

Gopalakrishnan V, Vignesh RC, Arunakaran J, Aruldhas MM, Srinivasan N (2006) Effects of glucose and its modulation by insulin and estradiol on BMSC differentiation into osteoblastic lineages. Biochem Cell Biol 84: 93–101

Guo Y, Xie C, Li X, Yang J, Yu T, Zhang R, Zhang T, Saxena D, Snyder M, Wu Y et al (2017) Succinate and its G-protein-coupled receptor stimulates osteoclastogenesis. Nat Commun 8: 15621

Hamann C, Goettsch C, Mettelsiefen J, Henkenjohann V, Rauner M, Hempel U, Bernhardt R, Fratzl-Zelman N, Roschger P, Rammelt S et al (2011) Delayed bone regeneration and low bone mass in a rat model of insulin-resistant type 2 diabetes mellitus is due to impaired osteoblast function. Am J Physiol Endocrinol Metab 301: E1220–1228

Hao ZC, Lu J, Wang SZ, Wu H, Zhang YT, Xu SG (2017) Stem cell-derived exosomes: A promising strategy for fracture healing. Cell Prolif 50

Hu Z, Ma C, Liang Y, Zou S, Liu X (2019) Osteoclasts in bone regeneration under type 2 diabetes mellitus. Acta Biomater 84: 402–413

Jeppesen DK, Fenix AM, Franklin JL, Higginbotham JN, Zhang Q, Zimmerman LJ, Liebler DC, Ping J, Liu Q, Evans R et al (2019) Reassessment of Exosome Composition. Cell 177: 428–445 e418

Jiao H, Xiao E, Graves DT (2015) Diabetes and Its Effect on Bone and Fracture Healing. Curr Osteoporos Rep 13: 327–335

Kishore R, Khan M (2017) Cardiac cell-derived exosomes: changing face of regenerative biology. Eur Heart J 38: 212–215

Kourembanas S (2015) Exosomes: vehicles of intercellular signaling, biomarkers, and vectors of cell therapy. Annu Rev Physiol 77: 13–27

Kume S, Kato S, Yamagishi S, Inagaki Y, Ueda S, Arima N, Okawa T, Kojiro M, Nagata K (2005) Advanced glycation end-products attenuate human mesenchymal stem cells and prevent cognate differentiation into adipose tissue, cartilage, and bone. J Bone Miner Res 20: 1647–1658

Li D, Liu J, Guo B, Liang C, Dang L, Lu C, He X, Cheung HY, Xu L, Lu C et al (2016) Osteoclast-derived exosomal miR-214-3p inhibits osteoblastic bone formation. Nat Commun 7: 10872

Li Z, Hassan MQ, Volinia S, van Wijnen AJ, Stein JL, Croce CM, Lian JB, Stein GS (2008) A microRNA signature for a BMP2-induced osteoblast lineage commitment program. Proc Natl Acad Sci U S A 105: 13906–13911

Liu X, Li Q, Niu X, Hu B, Chen S, Song W, Ding J, Zhang C, Wang Y (2017a) Exosomes Secreted from Human-Induced Pluripotent Stem Cell-Derived Mesenchymal Stem Cells Prevent Osteonecrosis of the Femoral Head by Promoting Angiogenesis. Int J Biol Sci 13: 232–244

Liu X, Yang Y, Li Y, Niu X, Zhao B, Wang Y, Bao C, Xie Z, Lin Q, Zhu L (2017b) Integration of stem cell-derived exosomes with in situ hydrogel glue as a promising tissue patch for articular cartilage regeneration. Nanoscale 9: 4430–4438

Marin C, Luyten FP, Van der Schueren B, Kerckhofs G, Vandamme K (2018) The Impact of Type 2 Diabetes on Bone Fracture Healing. Front Endocrinol (Lausanne) 9: 6

Min Z, Zhang R, Yao J, Jiang C, Guo Y, Cong F, Wang W, Tian J, Zhong N, Sun J et al (2015) MicroRNAs associated with osteoarthritis differently expressed in bone matrix gelatin (BMG) rat model. Int J Clin Exp Med 8: 1009–1017

Negishi-Koga T, Shinohara M, Komatsu N, Bito H, Kodama T, Friedel RH, Takayanagi H (2011) Suppression of bone formation by osteoclastic expression of semaphorin 4D. Nat Med 17: 1473–1480

Notsu M, Yamaguchi T, Okazaki K, Tanaka K, Ogawa N, Kanazawa I, Sugimoto T (2014) Advanced glycation end product 3 (AGE3) suppresses the mineralization of mouse stromal ST2 cells and human mesenchymal stem cells by increasing TGF-beta expression and secretion. Endocrinology 155: 2402–2410

Pando R, Even-Zohar N, Shtaif B, Edry L, Shomron N, Phillip M, Gat-Yablonski G (2012) MicroRNAs in the growth plate are responsive to nutritional cues: association between miR-140 and SIRT1. J Nutr Biochem 23: 1474–1481

Panganiban RP, Wang Y, Howrylak J, Chinchilli VM, Craig TJ, August A, Ishmael FT (2016) Circulating microRNAs as biomarkers in patients with allergic rhinitis and asthma. J Allergy Clin Immunol 137: 1423–1432

Paschou SA, Dede AD, Anagnostis PG, Vryonidou A, Morganstein D, Goulis DG (2017) Type 2 Diabetes and Osteoporosis: A Guide to Optimal Management. J Clin Endocrinol Metab 102: 3621–3634

Peinado H, Aleckovic M, Lavotshkin S, Matei I, Costa-Silva B, Moreno-Bueno G, Hergueta-Redondo M, Williams C, Garcia-Santos G, Ghajar C et al (2012) Melanoma exosomes educate bone marrow progenitor cells toward a pro-metastatic phenotype through MET. Nat Med 18: 883–891

Qin Y, Wang L, Gao Z, Chen G, Zhang C (2016) Bone marrow stromal/stem cell-derived extracellular vesicles regulate osteoblast activity and differentiation in vitro and promote bone regeneration in vivo. Sci Rep 6: 21961

Rauch A, Haakonsson AK, Madsen JGS, Larsen M, Forss I, Madsen MR, Van Hauwaert EL, Wiwie C, Jespersen NZ, Tencerova M et al (2019) Osteogenesis depends on commissioning of a network of stem cell transcription factors that act as repressors of adipogenesis. Nat Genet 51: 716–727

Sahoo S, Klychko E, Thorne T, Misener S, Schultz KM, Millay M, Ito A, Liu T, Kamide C, Agrawal H et al (2011) Exosomes from human CD34(+) stem cells mediate their proangiogenic paracrine activity. Circ Res 109: 724–728

Salazar-Roa M, Trakala M, Alvarez-Fernandez M, Valdes-Mora F, Zhong C, Munoz J, Yu Y, Peters TJ, Grana-Castro O, Serrano R et al (2020) Transient exposure to miR-203 enhances the differentiation capacity of established pluripotent stem cells. EMBO J 39: e104324

Schaap-Oziemlak AM, Raymakers RA, Bergevoet SM, Gilissen C, Jansen BJ, Adema GJ, Kogler G, le Sage C, Agami R, van der Reijden BA et al (2010) MicroRNA hsa-miR-135b regulates mineralization in osteogenic differentiation of human unrestricted somatic stem cells. Stem Cells Dev 19: 877–885

Su T, Xiao Y, Xiao Y, Guo Q, Li C, Huang Y, Deng Q, Wen J, Zhou F, Luo XH (2019) Bone Marrow Mesenchymal Stem Cells-Derived Exosomal MiR-29b-3p Regulates Aging-Associated Insulin Resistance. ACS Nano 13: 2450–2462

Takahara S, Lee SY, Iwakura T, Oe K, Fukui T, Okumachi E, Waki T, Arakura M, Sakai Y, Nishida K et al (2018) Altered expression of microRNA during fracture healing in diabetic rats. Bone & Joint Research 7: 139–147

Terpos E, Ntanasis-Stathopoulos I, Christoulas D, Bagratuni T, Bakogeorgos M, Gavriatopoulou M, Eleutherakis-Papaiakovou E, Kanellias N, Kastritis E, Dimopoulos MA (2018) Semaphorin 4D correlates with increased bone resorption, hypercalcemia, and disease stage in newly diagnosed patients with multiple myeloma. Blood Cancer J 8: 42

Toh WS, Lai RC, Hui JHP, Lim SK (2017) MSC exosome as a cell-free MSC therapy for cartilage regeneration: Implications for osteoarthritis treatment. Semin Cell Dev Biol 67: 56–64

Valadi H, Ekstrom K, Bossios A, Sjostrand M, Lee JJ, Lotvall JO (2007) Exosome-mediated transfer of mRNAs and microRNAs is a novel mechanism of genetic exchange between cells. Nat Cell Biol 9: 654–659

Waki T, Lee SY, Niikura T, Iwakura T, Dogaki Y, Okumachi E, Oe K, Kuroda R, Kurosaka M (2016) Profiling microRNA expression during fracture healing. BMC Musculoskelet Disord 17: 83

Wang N, Liu X, Shi L, Liu Y, Guo S, Liu W, Li X, Meng J, Ma X, Guo Z (2020) Identification of a prolonged action molecular GLP-1R agonist for the treatment of femoral defects. Biomater Sci 8: 1604–1614

Wu J, Kuang L, Chen C, Yang J, Zeng WN, Li T, Chen H, Huang S, Fu Z, Li J et al (2019) miR-100-5p-abundant exosomes derived from infrapatellar fat pad MSCs protect articular cartilage and ameliorate gait abnormalities via inhibition of mTOR in osteoarthritis. Biomaterials 206: 87–100

Xie Y, Chen Y, Zhang L, Ge W, Tang P (2017) The roles of bone-derived exosomes and exosomal microRNAs in regulating bone remodelling. J Cell Mol Med 21: 1033–1041

Xu G, Liu K, Anderson J, Patrene K, Lentzsch S, Roodman GD, Ouyang H (2012) Expression of XBP1s in bone marrow stromal cells is critical for myeloma cell growth and osteoclast formation. Blood 119: 4205–4214

Xu R, Shen X, Si Y, Fu Y, Zhu W, Xiao T, Fu Z, Zhang P, Cheng J, Jiang H (2018) MicroRNA-31a-5p from aging BMSCs links bone formation and resorption in the aged bone marrow microenvironment. Aging Cell 17: e12794

Yang K, Wang XQ, He YS, Lu L, Chen QJ, Liu J, Shen WF (2010) Advanced glycation end products induce chemokine/cytokine production via activation of p38 pathway and inhibit proliferation and migration of bone marrow mesenchymal stem cells. Cardiovasc Diabetol 9: 66

Ying W, Gao H, Dos Reis FCG, Bandyopadhyay G, Ofrecio JM, Luo Z, Ji Y, Jin Z, Ly C, Olefsky JM (2021) MiR-690, an exosomal-derived miRNA from M2-polarized macrophages, improves insulin sensitivity in obese mice. Cell Metab 33: 781–790 e785

Ying W, Riopel M, Bandyopadhyay G, Dong Y, Birmingham A, Seo JB, Ofrecio JM, Wollam J, Hernandez-Carretero A, Fu W et al (2017) Adipose Tissue Macrophage-Derived Exosomal miRNAs Can Modulate In Vivo and In Vitro Insulin Sensitivity. Cell 171: 372–384 e312

Zhang Y, Xie RL, Croce CM, Stein JL, Lian JB, van Wijnen AJ, Stein GS (2011) A program of microRNAs controls osteogenic lineage progression by targeting transcription factor Runx2. Proc Natl Acad Sci U S A 108: 9863–9868

Zhu Y, Jia Y, Wang Y, Xu J, Chai Y (2019) Impaired Bone Regenerative Effect of Exosomes Derived from Bone Marrow Mesenchymal Stem Cells in Type 1 Diabetes. Stem Cells Transl Med 8: 593–605

